# On minimising tumoural growth under treatment resistance

**DOI:** 10.1101/2023.07.04.547696

**Authors:** Matthias M. Fischer, Nils Blüthgen

## Abstract

Drug resistance is a major challenge for curative cancer treatment, representing the main reason of death in patients. Evolutionary biology suggests pauses between treatment rounds as a way to delay or even avoid resistance emergence. Indeed, this approach has already shown promising preclinical and early clinical results, and stimulated the development of mathematical models for finding optimal treatment protocols. Due to their complexity, however, these models do not lend themself to a rigorous mathematical analysis, hence so far clinical recommendations generally relied on numerical simulations and ad-hoc heuristics. Here, we derive two mathematical models describing tumour growth under genetic and epigenetic treatment resistance, respectively, which are simple enough for a complete analytical investigation. First, we find key differences in response to treatment protocols between the two modes of resistance. Second, we identify the optimal treatment protocol which leads to the largest possible tumour shrinkage rate. Third, we fit the ”epigenetic model” to previously published xenograft experiment data, finding excellent agreement, underscoring the biological validity of our approach. Finally, we use the fitted model to calculate the optimal treatment protocol for this specific experiment, which we demonstrate to cause curative treatment, making it superior to previous approaches which generally aimed at stabilising tumour burden. Overall, our approach underscores the usefulness of simple mathematical models and their analytical examination, and we anticipate our findings to guide future preclinical and, ultimately, clinical research in optimising treatment regimes.

## 1. Introduction

Despite tremendous advances in the field of cancer research, resistance to treatment has remained a major challenge for the curative eradication of tumours and is considered the main reason of death in cancer patients (Vasan et al., 2019). Accumulating evidence strongly suggests intratumoural heterogeneity, i.e. tumours being a mixture of functionally distinct cancer cells differing either genetically or epigenetically, as a major driver of drug resistance (Saunders et al., 2012; Pribluda et al., 2015). Under the selective pressure exerted by treatments, resistance can then emerge as a result of either the expansion of pre-existing resistant subpopulations or the emergence of new drug-tolerant cells (Dagogo-Jack and Shaw, 2018).

From a perspective of evolutionary biology, it thus appears promising to reduce such a selective pressure towards resistance by introducing strategically placed pause phases into treatment protocols, during which no treatment is applied. Indeed, such ”evolutionary strategies” have already shown promise in preclinical and early-stage clinical trials (Silva et al., 2012; Enriquez-Navas et al., 2016; Zhang et al., 2017). They have also stimulated the development and analysis of mathematical models aiming to capture the essence of these processes and ultimately find optimal treatment protocols (Smalley et al., 2019; Kim et al., 2021). While describing experimental data with a high degree of accuracy and having a demonstrably large amount of predictive power, these models do not lend themself to a rigorous mathematical analysis due to the involved nonlinear terms which make deriving closed-form solutions of the occuring dynamics impossible.

To close this gap in the literature, in this work we derive and study two mathematical models describing the population dynamics of a growing tumour for the cases of a genetically and an epigenetically acquired treatment resistance, respectively. Importantly, both models are simple enough to be entirely accessible to analytical investigation. First, we compare the dynamics of these two models, finding key differences in growth and response to treatment protocols. Subsequently, we identify and characterise the optimal treatment protocol which leads to the asymptotically smallest tumour growth, i.e. the largest possible rate of tumour shrinkage. Afterwards, we fit our model of phenotypically acquired treatment resistance to previously published experimental data from mouse experiments with patient-derived melanoma xenografts harbouring acquired resistances to *MEK* -inhibitor treatment (Hong et al., 2018). Doing so, we find excellent agreement between data and theory, which underscores the predictive power of our modelling approach. Finally, we use the fitted model to calculate the optimal treatment protocol for this specific experiment and perform numerical simulations to demonstrate that the obtained solution is indeed able to drive the tumour into complete extinction and hence outperforms earlier ad-hoc approaches.

## 2. Materials and Methods

### 2.1. Genetic model

We assume a tumour consisting of two different cell types *S, R*, denoting treatment-sensitive and treatment-resistant cells, respectively. During treatment, the compartments grow exponentially at rates 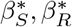, respectively, whereas in the absence of treatment their growth rates amount to *β*_*S*_, *β*_*R*_, respectively. We decided to not introduce a carrying capacity because we assume that the overall tumour size is sufficiently small compared to the maximum possible tumour size, and thus growth to not be limited yet. We will revisit this assumption later. Resistance to treatment emerges due to genetic mutations at a rate *µ*, which is small enough for exact back-mutations to a sensitive cell state to be neglectable. Thus, during treatment the system is governed by the following set of differential equations

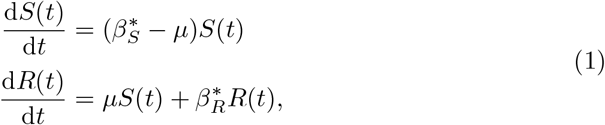

and during treatment pauses by

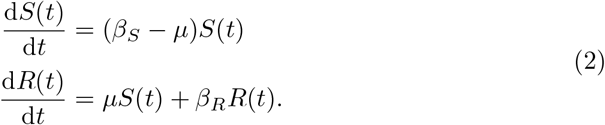

### 2.2. Phenotypic plasticity model

The ”phenotypic plasticity model” is similar to the genetic model described in the previous section. However, we assume that resistance emerges as a consequence of treatment at a constant rate *α*, whereas in the absence of treatment resistance is lost again at a rate *δ*. Thus, during treatment the system is governed by the following set of differential equations

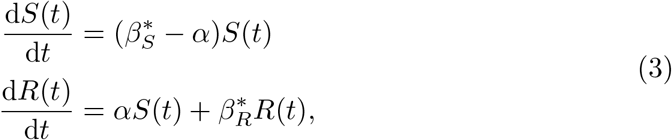

and during treatment pauses by

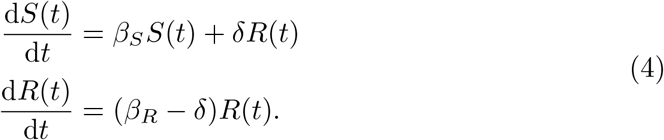

### 2.3. Literature data, model fitting and illustrative model simulation

Tumour size time-course data from mouse xenograft experiments have been extracted from Figures 5a and 5b of Hong et al. (2018) using the WebPlotDigitizer software (https://automeris.io/WebPlotDigitizer/), which has been found to be a reliable and valid tool for extracting data from diagrams (Drevon et al., 2017). Model fitting was performed using the `optimize.curve_fit` routine from the `scipy` package (Virtanen et al., 2020), version 1.10.1, for the Python programming language (Van Rossum and Drake Jr, 1995), version 3.9.6. The routine implements unconstrained non-linear least square optimisation via the Levenberg-Marquardt algorithm. For illustrative purposes, the model has been simulated numerically using the `integrate.solve_ivp` routine from the same software package, which relies on an explicit Runge-Kutta method with a fourth-order error control method and a step size selection using a fifth-order accurate formula with local extrapolation. All computations have been carried out on a 2023 MacBook Pro with an Apple M2 Max processor running macOS Ventura, version 13.3.

## 3. Results

### 3.1. Phenotypic plasticity requires a precisely timed treatment protocol

We derived two simple mathematical models of tumoural resistance to treatment, both of which distinguish between compartments *S* and *R*, denoting sensitive and resistant cells, respectively. In both models, during treatment the two compartments grow at rates 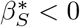 and 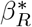, respectively. During treatment pauses, they grow at different rates *β*_*S*_ *>* 0, *β*_*R*_, respectively.

In the ”genetic model”, treatment resistance is acquired via random mutations at a small rate *µ* regardless of whether a treatment is applied or not. Due to *µ* being small, we neglect the possibility of back-mutations. In contrast, in the ”phenotypic plasticity model”, resistance is induced at a rate *α* during treatment phases, and lost again at a rate *δ* during treatment pauses. For an illustration refer to Figure 1A and Figure 1B.

**Figure 1.**
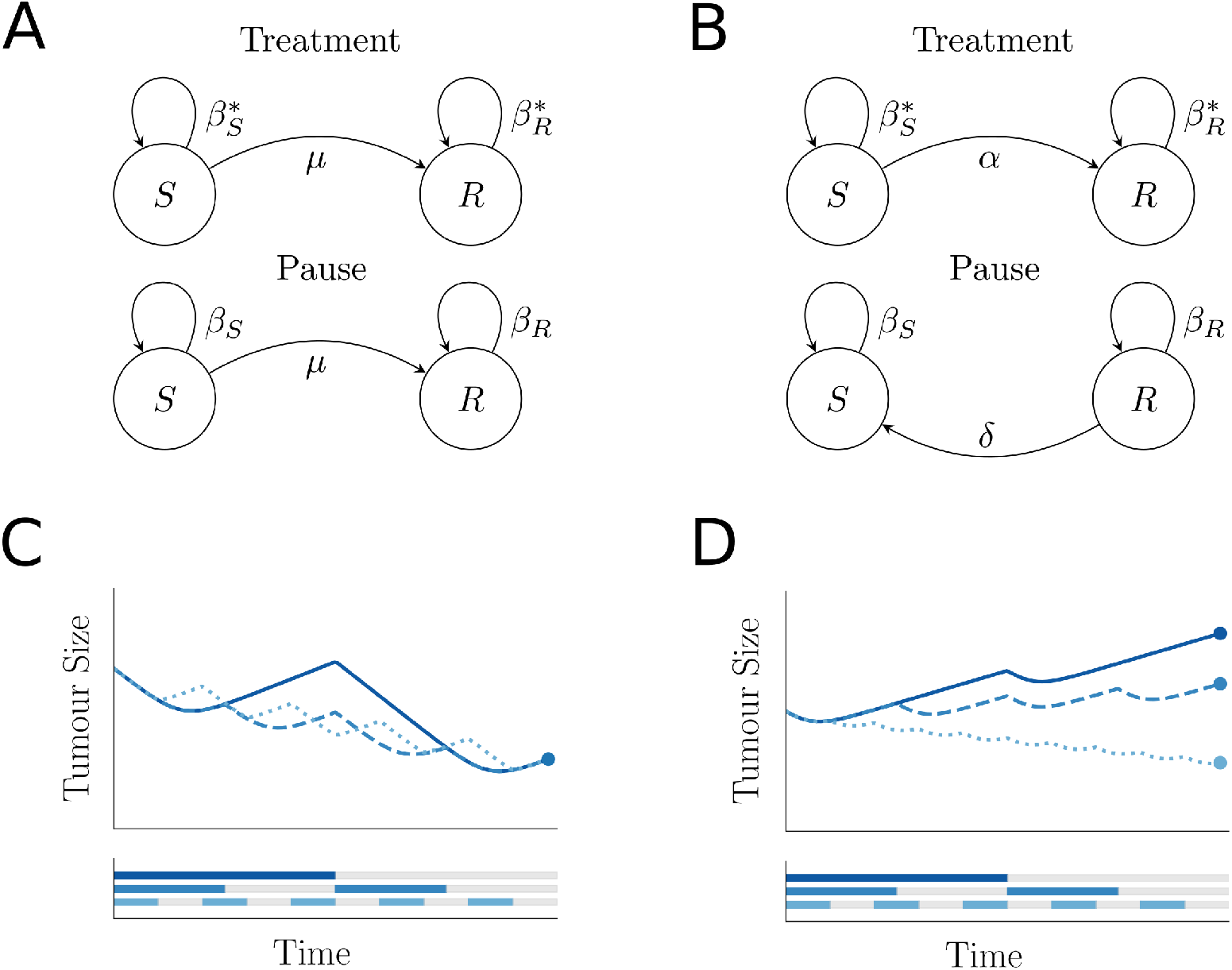
Phenotypic plasticity requires a precisely timed treatment protocol. **A** Schematic sketch of the model with treatment resistance emerging from random mutations (”genetic model”) at a small rate *µ. S*: Sensitive subpopulation, *R*: Resistant subpopulation 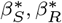 denote the growth rates of the compartments during treatment; *β*_*S*_, *β*_*R*_ during treatment pause. **B** Schematic sketch of the model with acquired treatment resistance due to phenotypic plasticity (”phenotypic plasticity model”). Growth rates are the same as before, however during treatment sensitive cells acquire resistance at a rate *α*, and in the absence of treatment resistant cells lose resistance at a rate *δ*. **C** Final tumour size in the genetic model is not affected by the ordering of the individual treatment and pause phases. Numerical illustration. **D** In the phenotypic model, the order of the individual treatment and pause phases affects the final tumour size. Numerical illustration.

For both models, we define *y*(*t*) := [*S*(*t*), *R*(*t*)]^*T*^ and can write the system as

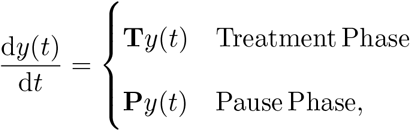

where we have in case of the genetic model (Equations 1 and 2)

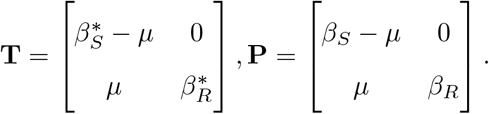

and in case of the phenotypic plasticity model (Equations 3 and 4).

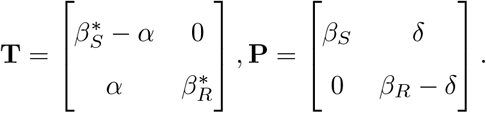

We define a treatment protocol *𝒫* as a sequence of alternating treatment and pause phases. We write *𝒫* = (*τ*_1_, *φ*_1_, *τ*_2_, *φ*_2_, …, *τ*_*i*_, *φ*_*i*_), where *τ*_*i*_ represents the length of the *i*-th treatment phase, and *φ*_*i*_ the length of the *i*-th pause phase.

For the genetic model (Equations 1 and 2), we assume that the mutation rate *µ* is sufficiently small to be neglected. This decouples the two system compartments, and thus the final compartment sizes *S*_*𝒫*_, *R*_*𝒫*_ can be directly calculated as a product of exponential growth terms:

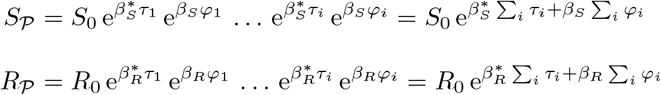

Thus, given any initial condition (*S*_0_, *R*_0_) and set of system parameters 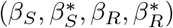, only the overall sum 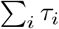 of treatment phase lengths and the overall sum 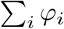 of pause phase lengths determine the final tumour size, but not their segmentation or their order. We illustrate this finding in Figure 1C.

In contrast, for the phenotypic plasticity model (Equations 3 and 4), the final system state *y*_*𝒫*_ = [*S*_*𝒫*_, *R*_*𝒫*_]^*T*^ is given by

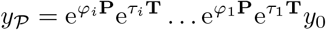

Since all matrix exponentials 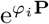 and 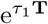 are upper and lower triangular matrices, respectively, they do not commute in general. Hence, the order of treatment and pause phases will influence the final tumour size. We illustrate this finding in Figure 1D.

Taken together, from a biological point of view, these results show that the growth, and thus the successful eradication of tumours with exclusively genetic resistance mechanisms solely depend on the sum of time spent in treatment vs. in pause phase. Conversely, the order of the individual phases of a treatment regimen does not influence the final outcome. In contrast, tumours which show an acquired treatment resistance stemming from phenotypic plasticity demand a precise segmentation and ordering of the individual treatment and pause phases to be optimally controlled.

### 3.2. Change of variables identifies the optimal treatment protocol

In a clinical setting, usually we can only monitor the overall tumour burden, for instance by using medical imaging or by measuring the concentration of markers of overall tumour burden. In contrast, generally we are not able to track the number of sensitive vs. resistant cells. This motivates us to perform the following change of variables: Let *N* (*t*) := *R*(*t*) + *S*(*t*) denote the observable overall tumour burden, and *Q*(*t*) := *R*(*t*)*/*(*R*(*t*) + *S*(*t*)) denote the ”invisbile” fraction of resistant cells present in the tumour (see Figure 2A for an illustration).

**Figure 2.**
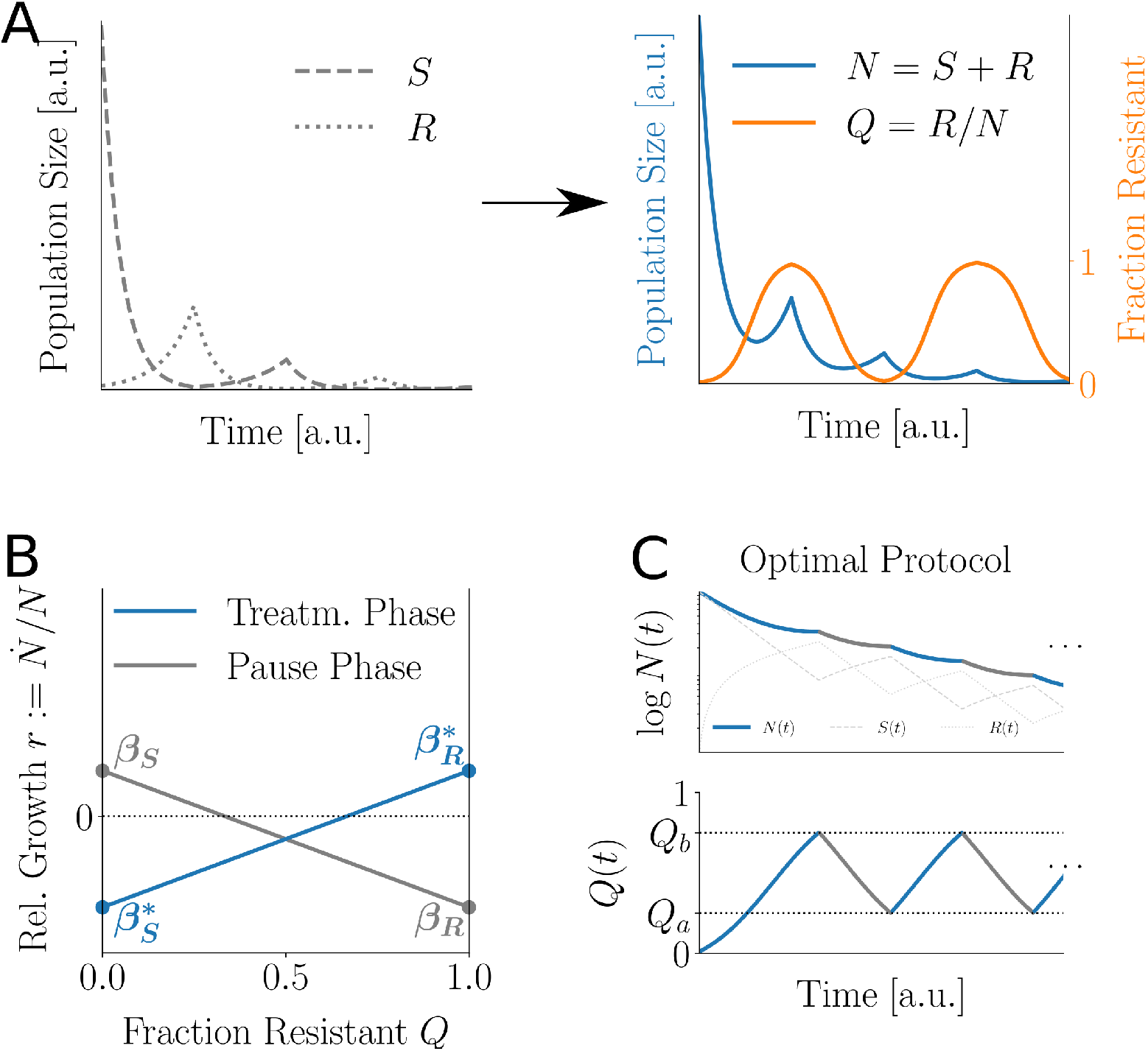
Rewriting the models in terms of overall population size *N* and fraction *Q ∈* [0; 1] of resistant cells reveals optimal treatment protocol. **A** Illustration of the model transformation. **B** In both the treatment phase (blue line) and the pause phase (grey line), relative local tumour growth 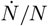 is a simple linear function of *Q*, determined by the subpopulation growth rates. **C** Illustration of the analytically derived optimal treatment protocol: After a lead-in treatment, during which *Q* reaches some *Qa*, we repeatedly switch between treatment (until *Q* reaches some *Q*_*b*_ *> Q*_*a*_), and treatment pause (during which *Q* recedes back to *Q*_*a*_). Upper diagram shows log population size over time (continuous blue line during treatment, continuous grey line during pause). Additionally, the sizes of the sensitive and resistant subpopulation are plotted with a dashed and a dotted light grey line, respectively. Lower diagram shows fraction *Q* of resistant cells over time.

Both models then satisfy

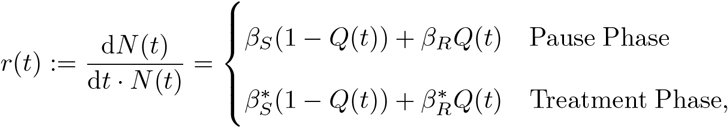

which, importantly, shows that in both phases the overall relative tumour growth *r*(*t*) is only a linear function of the tumour composition given by *Q*(*t*) (see Figure 2B for an illustration).

It can easily be verified that during treatment *Q* follows a monotonically increasing sigmoid function converging to unity, whereas during treatment pauses, *Q* follows a monotonically decreasing function which converges to zero. Assuming that a treatment-naive tumour mostly consists of sensitive cells, the relative tumour growth rate before any treatment is approximately identical to *β*_*S*_, and the relative growth rate directly after the onset of treatment approximately equals 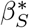. During treatment, *r*(*t*) will tend towards 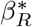. Conversely, during treatment pause, *r*(*t*) will tend to *β*_*S*_ again. It should be noted that these results also imply that it is possible to infer the current tumour composition by using sufficiently frequent measurements of overall tumour burden alone. In particular, for the case of phenotypic plasticity knowledge of the transition rates *α, δ* is not required.

We now want to find the optimal treatment protocol, defined as the treatment protocol which, after transient behaviour has passed, leads to minimal relative tumour growth *r* per unit of time. From the results of the previous section it follows that any treatment protocol corresponds to a continuous trajectory through *Q*. As per Equation 3.2, for any such trajectory *θ*, there exists a corresponding piecewise continuous function *r*_*θ*_, describing the local growth rate of *N* over time. Accordingly, the final tumour population *N*_*θ*_ at the end of *θ* is given by

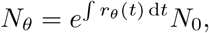

where the integral has to be broken into the individual treatment and pause phases. Obviously, every finite trajectory through *Q* will only lead to a finite relative reduction in tumour size, as the exponential term is strictly greater than zero. Hence, we wish to find a closed path through *Q* which we can follow over and over again in order to force *N* (*t*) *→* 0.

In order to further characterise the optimal solution, we can prove (see Appendix A) that among all closed trajectories, a trajectory can only be optimal if it is free of any inner loops. In other words, the closed trajectory which minimises relative tumour growth per unit of time consists of one treatment phase, where *Q*(*t*) grows from *Q*_*a*_ to *Q*_*b*_, followed by one pause phase, where *Q*(*t*) decreases back to *Q*_*a*_.

Accordingly, given any system parametrisation, it suffices to find the optimal switching points *Q*_*a*_, *Q*_*b*_, and the optimal treatment protocol consists of initial transient phase where *Q*(*t*) approaches *Q*_*a*_ from its initial value, followed by repeatedly applying treatment until *Q*_*b*_ is reached and pausing treatment, until *Q*_*a*_ is reached again (see Figure 2C for an illustration).

### 3.3. Our approach accurately describes literature data and predicts the optimal treatment schedule

After examining the models analytically, we fitted the phenotypic model to time-course tumour burden data from mouse xenograft experiments published in Hong et al. (2018). First, we used the data from Figure 5a, where patient derived melanoma xenografts sensitive to *MEK* -inhibitor treatment were implanted in mice and grown for 18 days. Afterwards, treatment was started by giving 5 mg/kg *MEK* -inhibitor daily for additional 67 days. Because tumour growth started to saturate and became subexponential around day 40, we discarded all the data points from this point of time. This time-course describes the growth of sensitive tumour cells *in vivo*, their reaction to the onset of treatment, and the acquisition of treatment resistance (see Figure 3A, left panel).

**Figure 3.**
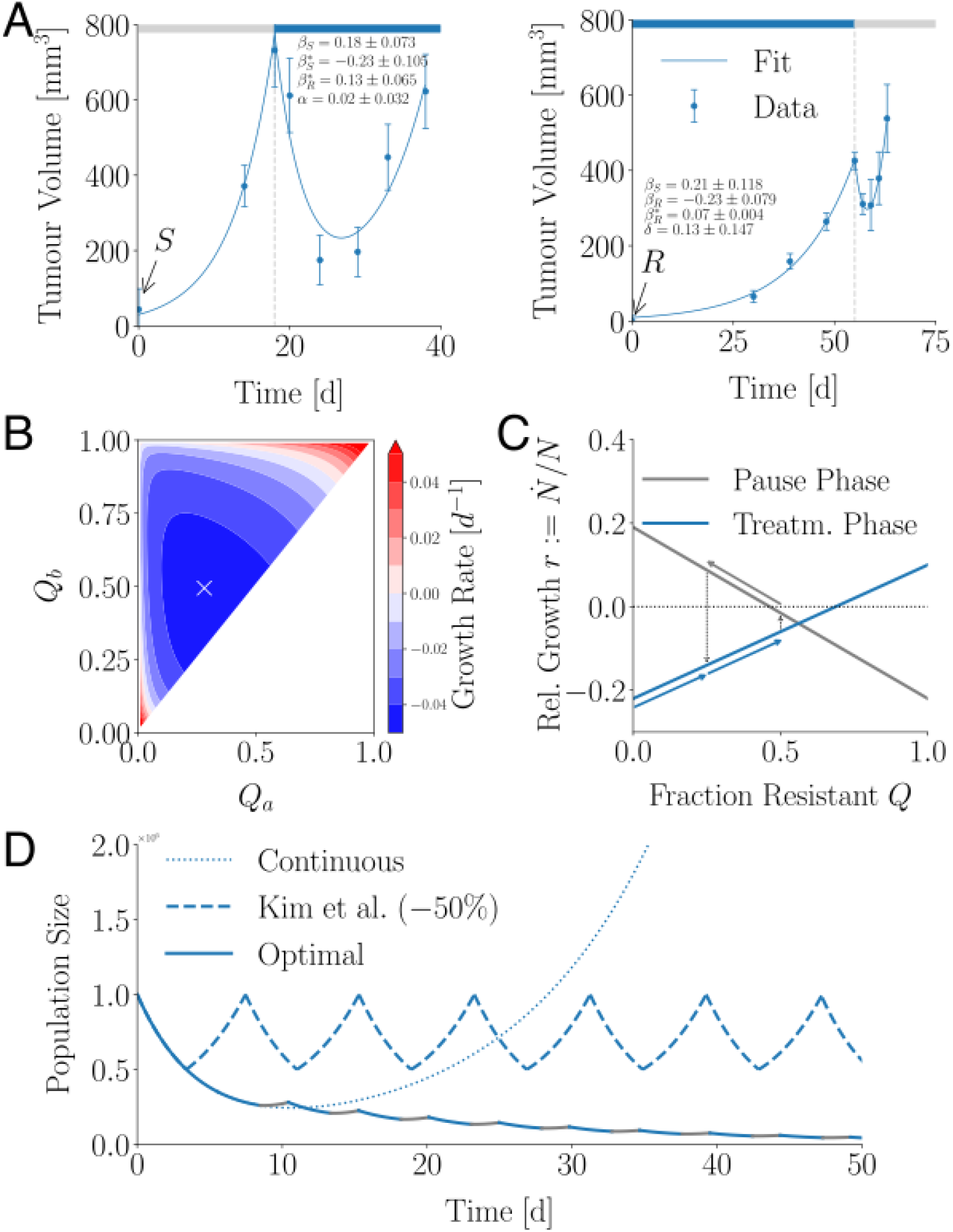
Application of the phenotypic model to the data by Hong et al. (2018). **A** Left: Tumour xenograft volume data (blue dots), model fit (continuous blue lines) and obtained parameter estimates for the data from Fig 5A, sensitive population subjected to MEKi treatment. Dashed grey line denotes the onset of treatment. Right: The same for Figure 5B, resistant cells subjected to MEKi withdrawal. Dashed grey line denotes the termination of treatment. Observe the very good fit for both data sets and the agreement between shared parameters. **B** Calculated tumour growth rate per day as a function of chosen lower and upper switch points *Q*_*a*_, *Q*_*b*_. Cross marks *Q*_*a*_ = 0.25, *Q*_*b*_ = 0.5 (i.e. 25% and 50% resistant cells), which will be examined in the rest of this figure. **C** Relative local tumour growth rate 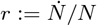 as a function of resistance fraction *Q* for pause (continuous grey line) and treatment phase (continuous blue line). Arrows illustrate the treatment protocol with switch points as chosen before. **D** Numerical simulation of a tumour population over time treated as defined before (continuous blue line during treatment, continuous grey line during pause phase). Note the convergence to tumour extinction, whereas continuous treatment (dotted blue line) would cause relapse after ca. 10 days, and the ”50% stopping rule” by Kim et al. (2021) (dashed blue line) would only lead to a stabilisation of tumour burden. Note that the proposed ”20% stopping rule” from the same paper would not work here, as continuous treatment cannot achieve a tumour reduction by 80%.

Second, we also fitted the model to data from Figure 5b of the same paper, where the same cell line was first made resistant to treatment by subjecting them to inhibitor treatment *in vitro*. Resistant cells were then implanted into mice, which were treated with 5 mg/kg *MEK* -inhibitor daily for 52 days, after which treatment was stopped for additional 23 days. Accordingly, this time-course describes the growth of resistant cancer cells during treatment, their reaction to the termination of treatment, and the subsequent loss of resistance and re-emergence of the sensitive phenotype (see Figure 3A, right panel). Note the excellent fit of the model to the data in both cases, as well as very similar estimates of the two shared parameters across the two fits.

After parametrisation, we used the model to find the optimal treatment protocol for this specific set of parameters, which means finding the optimal switching points *Q*_*a*_ *> Q*_*b*_ in fraction of resistant cells (see previous section). Hence, we calculated the realised tumour growth rate as a function of *Q*_*a*_ and *Q*_*b*_, which is depicted in Figure 3B, where blue colors denote areas of negative growth, i.e. a shrinking tumour. Obviously, as *Q*_*a*_ *→ Q*_*b*_, the duration of treatment cycles shrinks, making the realised growth rate diverge to *−∞*. However, in reality treatment cycles obviously cannot become arbitrarily short. For this reason, we selected the scenario of *Q*_*a*_ = 0.25, *Q*_*b*_ = 0.5 (marked with a white cross in the figure) as a reasonable compromise between sufficient tumour shrinkage per day and feasibility in reality for further investigation.

Analogously to Figure 2B, Figure 3C shows the relative tumour growth *r* as a function of the fraction *Q* of resistant cells during treatment phase (blue line) and pause phase (grey line) for the obtained parametrisation. Little blue and grey arrows depict the treatment protocol we have selected in the previous paragraph, i.e. an initial lead-in treatment from *Q ≈* 0 to *Q*_*a*_ = 0.25, followed by repeatedly applying treatment until *Q* reaches *Q*_*b*_ = 0.5 and pausing treatment until *Q* has reached *Q*_*a*_ = 0.25 again.

Numerically simulating this treatment protocol indeed confirms that it will lead to tumour extinction (continuous line depicted in Figure 3D, where blue and grey colours indicates treatment and pause phase, respectively). in contrast, classical continuous treatment (dotted blue line) only causes an initial rediction in tumour size, followed by tumour relapse and the onset of exponential growth. We also simulated the ”50 % rule” mentioned and studied in Kim et al. (2021), where treatment is applied until the tumour has reached half of its initial size, and then paused until the tumour has reached its entire initial size again, where treatment is then resumed (dashed blue line). This approach succeeds in permanently stabilising tumour size, but cannot achieve curative treatment. Kim et al. (2021) also proposed the ”20% rule”, where treatment is applied until the tumour has shrunk to a fifth of its initial size instead. Of note, this rule would fail for this specific parametrisation, as continuous treatment (see dotted line) is not able to cause such a drastic reduction in the first place.

Taken together, we demostrated that, for the specific case of the experiments by Hong et al. (2018) our model describes the data well and can be used to identify an optimal treatment protocol that is superior to previously published approaches.

## 4. Discussion

In this work, we have derived and analysed two simple mathematical models describing tumoural resistance to treatment via genetic and epigenetic mechanisms, respectively. We have used these models to solve the problem of introducing treatment pauses to minimise tumour growth in the face of such resistances. In contrast to preceding modelling efforts, we here aimed for utmost simplicity, in particular by not including any growth limiting carrying capacities. While this simplifying assumption is of course a departure from biological reality, we still considered it necessary to build models accessible to a complete analytical examination, and thus be able to obtain reliable insights independently of specific parametrisations and potentially costly numerical simulations. The validity of this approach is demonstrated by the fact that despite this simplification it was still able to describe experimental data (Hong et al., 2018) with a high degree of accuracy.

Even without a carrying capacity, in our models it can be possible to achieve complete tumour eradication, which is conceptually intriguing. This, however, necessarily requires that resistant cells experience negative growth during treatment pauses. Such a ”drug addiction” of resistant cells has been observed repeatedly not only in cell culture (Suda et al., 2012; Sun et al., 2014; Moriceau et al., 2015) and animal models (Das Thakur et al., 2013), but also *in vivo* (Dietrich et al., 2013; Seifert et al., 2016; Dooley et al., 2016), and can thus be expected to be a viable option to exploit therapeutically in some patients. Nonetheless, even if during treatment pauses resistant cells grow only slower than sensitive cells but still at a positive rate, in our models intermittent treatment will still minimise overall tumour growth compared to continuous treatment protocols. All of these findings agree with numerical results from a more complex competitive Lotka-Volterra-like system (Strobl et al., 2021), in which additionally the role of cellular turnover was found to be an important predictor of therapy success as it amplifies the effects of competition and resistance costs. Accordingly, an interesting and straightforward extension of our work might be the explicit introduction of a cellular turnover rate (or ”apoptosis rate”) as opposed to using aggregated subpopulation net growth rates.

For the specific case of genetic resistances, we have shown that overall tumour growth is only determined by the sum of time spent in treatment vs. pause phase. In contrast, the exact segmentation and ordering of the individual treatment and pause phases does not influence overall tumour size at the end of a treatment protocol. From a clinical point of view, this implies that there is not much room for improvement in treatment protocols apart from adjusting the fraction of time spent in treatment pause. One could, however, argue that a relatively long initial lead-in treatment before any treatment pauses might be advantageous, as it would reduce the cumulative risk of resistance emergence via random mutations. Importantly, this holds only if it can be guaranteed that the treatment does not induce the acquisition of further resistance. As a direct consequence, this finding also underscores the need for elucidating the exact mechanisms responsible for a given tumour becoming resistant as these crucially affect the optimal treatment strategy (Housman et al., 2014).

In stark contrast to genetically conferred resistance, in case of epigenetic mechanisms, the order of the individual phases of a treatment protocol can profoundly influence tumour growth, and thus the possibility of curative tumour eradication. Hence, tumours harboring epigenetic resistances require a precisely tuned treatment protocol to maximise chances of complete tumour extinction.

The main contribution of our work solves this problem by reducing this problem to navigating along an one-dimensional axis which encodes the fraction of currently resistant cells in the tumour. In this way, it becomes apparent that the optimal protocol, which minimises overall tumour size, keep the fraction of resistant cells in a range [*Q*_*a*_, *Q*_*b*_] which can be calculated numerically. As shown with experimental data (Hong et al., 2018), our approach can in some cases even cause complete tumour eradication, which is a qualitative improvement over previous approaches such as Kim et al. (2021) which generally aimed for stabilising tumour burden in a pre-specified range. The drawback of our approach obviously lies in the fact that we need to know the parameter values describing the tumour of a given specific patient to be able to calculate the treatment protocol optimal for this specific patient. This, in turn, requires sufficiently frequent and accurate measurements of tumour burden over time, so that the model can be proeprly parametrised. In itself, this is not a particularly strong requirement and similar to previous approaches such as Smalley et al. (2019). Particlarly for leukemia, such a freqent monitoring of the tumor burden is relatively straight forward and minimally invasive. However, the need to perform curve-fitting on these data afterwards and interpret the obtained results is likely to pose a challenge in a clinical setting.

Nonetheless, if this challenge is solved and such a real-time analysis of tumour burden data is possible in the clinic, an idealised scenario of cancer treatment may look as follows. Upon initial admission of the patient, data is collected as sketched out in Figure 4. Before any treatment, tumour burden is measured multiple times, so that the growth rate of the treatment-naive and hence -sensitive tumour cells can be estimated (corresponding to the parameter *β*_*S*_ in our model). Directly after the onset of the first round of treatment, the tumour is still mostly comprised of sensitive cells, so its rate of shrinking corresponds to the growth rate of sensitive cells under treatment 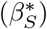. During treatment, sensitive cells continue to die, however they may also switch to a resistant cell state at a rate we cannot measure directly (*α*). Over time, the majority of tumour cells will have become resistant, and thus the tumour growth rate will converge to the growth rate of resistant cells under treatment 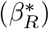 until at some point the relapse becomes obvious. Treatment is then halted, and directly afterwards the now mostly-resistant tumour approximately follows the growth rate of untreated resistant cells (*β*_*R*_, which is less than zero in case of drug addiction). Without treatment, however, resistant cells will also lose resistance at another rate we cannot directly measure (*δ*). Accordingly, the entirety of the collected data is now used for curve-fitting. Because the raw data already provide good estimates or bounds for the four growth rates, the remaining two parameters *α, δ* generally will be identifiable, as demonstrated in Section 3.3. Now that all relevant parameters are estimated, the obtained parameters are used to computationally identify the optimal course of treatment, which is initiated afterwards. This optimal treatment is guaranteed to minimise overall tumour growth and may, in some cases, even lead to curative treatment.

**Figure 4.**
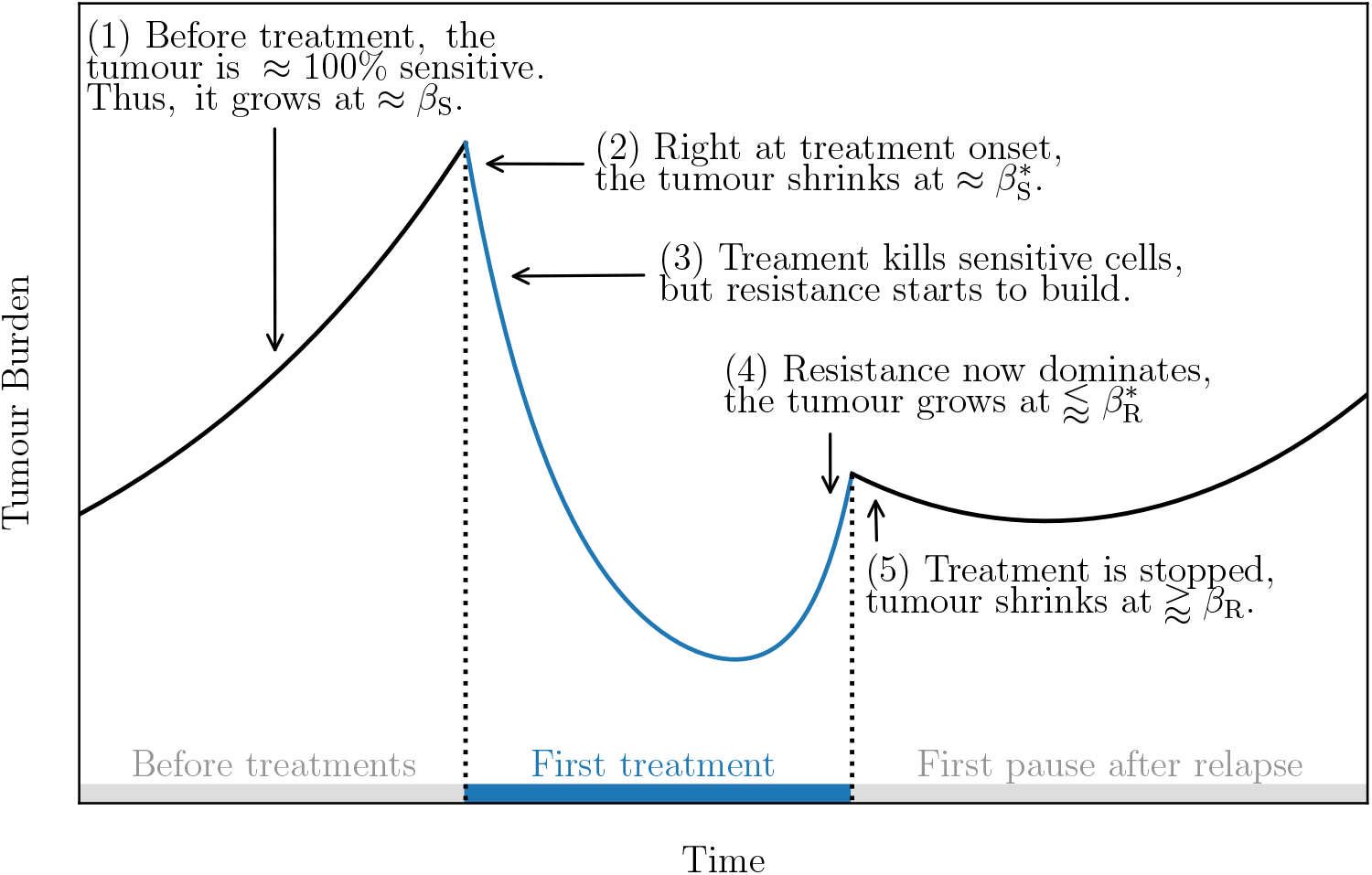
Data collection protocol for an idealised scenario of cancer treatment with real-time monitoring of tumour burden. **(1)** Before treatment, the tumour mostly consists of sensitive cells and thus follows the growth rate of untreated sensitive cells (*β*_*S*_ *>* 0). **(2)** Directly after treatment onset, the still sensitive tumour follows the growth rate of sensitive cells under treatment 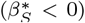. **(3)** During treatment, sensitive cells continue to die, however may also switch to a resistant state at rate *α*, which is not directly observable and needs to be identified via curve-fitting once all data have been acquired. **(4)** Once the majority of tumour cells has become resistant, the tumour follows the growth rate of resistant cells under treatment 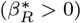. **(5)** Treatment is halted, and directly afterwards the now mostly-resistant tumour approximately follows the growth rate of untreated resistant cells (*β*_*R*_, which is less than zero in case of drug addiction). Without treatment, resistant cells will also lose resistance at a rate *δ* only identifiable via curve-fitting. **Not shown:** Once estimates of all six parameters have become available from curve-fitting, they are used to computationally identify the optimal course of treatment, which is initiated afterwards. This optimal treatment is guaranteed to minimise overall tumour growth and may, in some cases, even lead to curative treatment.

Our analysis shows that optimal treatment protocols may differ for every individual patient. This is in agreement with a general tendency against a ”one-size-fits-all approach” in medicine and argues for true ”precision medicine” or ”personalised medicine” where treatment is individually taylored to every given patient (King and Robins, 2006). In contrast to the majority of approaches in precision medicine (König et al., 2017), however, we here did not rely on any molecular data such as genomic, transcriptomic, or proteomic profiling. Instead, we personalised our model by fitting it to solely phenotypical data, i.e. tumour burden measurements over time. In addition, instead of using involved and potentially hard to interpret computational pipelines, our modelling approach aimed for utmost simplicity and its success underscores the usefulness of simple mathematical models and their thorough analytical examination. Thus, we anticipate not only our immediate results, but also our general methodological approach to guide and inspire future preclinical and, ultimately, clinical research in precision medicine in general, and optimising cancer treatment regimes in particular.

## 5. Acknowledgements

MMF would like to thank Rosario Astaburuaga-García for helpful comments on the data analytic part of this work, and acknowledges funding by the DFG (Deutsche Forschungsgemeinschaft, RTG2424, CompCancer).

## Declarations

### 5.1. Author contributions

MMF carried out model derivations and analyses with help from NB, and wrote the initial version of the manuscript. NB contributed to the final version of the manuscript and supervised the project. All authors have read and approve of the final version of this manuscript.

### 5.3. Code availability

The complete commented source code used for all numerical examinations presented in this manuscript are available in the following github repository: https://github.com/Matthias-M-Fischer/Minimising.

## Appendix A. Optimality of trajectories without inner loops

First, we define a closed trajectory *π* through *Q* to be *optimal*, exactly if the relative reduction of tumour size it causes per unit of time is greater than or equal to all possible other closed trajectories through *Q*.

Now, we prove the following

### Proposition 1.

*A closed trajectory through Q can only be optimal if it is free of any inner loops*.

*Proof*. Assume the opposite, that is, the optimal path *π* through *Q* consists of *n* inner loops. Due to the commutativity of products of exponentials (see Equation 3.2), the final tumour volume does not change if we rearrange *π* into *n* + 1 separate closed ”atomic” loops *π*_1_ … *π*_*n*+1_ without inner loops. Starting with an initial tumour size *N*_0_, one full traversal of the *i*-th loop takes 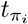 units of time, leading to a relative tumour reduction of 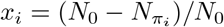. Let *π*^*\**^ denote the inner loop which maximises *x*_*i*_*/t*_*i*_. Then, however, it is easy to show that *π*^*\**^ reduces tumour growth more efficiently than *π*. Contradiction. □

## References

Dagogo-Jack, I. and Shaw, A. T. (2018). Tumour heterogeneity and resistance to cancer therapies. Nature reviews Clinical oncology, 15(2):81–94.

Das Thakur, M., Salangsang, F., Landman, A. S., Sellers, W. R., Pryer, N. K., Levesque, M. P., Dummer, R., McMahon, M., and Stuart, D. D. (2013). Modelling vemurafenib resistance in melanoma reveals a strategy to forestall drug resistance. Nature, 494(7436):251–255.

Dietrich, S., Hüllein, J., Hundemer, M., Lehners, N., Jethwa, A., Capper, D., Acker, T., Garvalov, B. K., Andrulis, M., Blume, C., et al. (2013). Continued response off treatment after braf inhibition in refractory hairy cell leukemia. Journal of clinical oncology: official journal of the American Society of Clinical Oncology, 31(19):e300–3.

Dooley, A. J., Gupta, A., and Middleton, M. R. (2016). Ongoing response in braf v600e-mutant melanoma after cessation of intermittent vemurafenib therapy: a case report. Targeted oncology, 11:557–563.

Drevon, D., Fursa, S. R., and Malcolm, A. L. (2017). Intercoder reliability and validity of webplotdigitizer in extracting graphed data. Behavior modification, 41(2):323–339.

Enriquez-Navas, P. M., Kam, Y., Das, T., Hassan, S., Silva, A., Foroutan, P., Ruiz, E., Martinez, G., Minton, S., Gillies, R. J., et al. (2016). Exploiting evolutionary principles to prolong tumor control in preclinical models of breast cancer. Science translational medicine, 8(327):327ra24–327ra24.

Hong, A., Moriceau, G., Sun, L., Lomeli, S., Piva, M., Damoiseaux, R., Holmen, S. L., Sharpless, N. E., Hugo, W., and Lo, R. S. (2018). Exploiting drug addiction mechanisms to select against mapki-resistant melanomaa synthetic lethality underlying mapki addiction in melanoma. Cancer Discovery, 8(1):74–93.

Housman, G., Byler, S., Heerboth, S., Lapinska, K., Longacre, M., Snyder, N., and Sarkar, S. (2014). Drug resistance in cancer: an overview. Cancers, 6(3):1769–1792.

Kim, E., Brown, J. S., Eroglu, Z., and Anderson, A. R. (2021). Adaptive therapy for metastatic melanoma: Predictions from patient calibrated mathematical models. Cancers, 13(4):823.

King, R. J. B. and Robins, M. W. (2006). Cancer biology. Pearson Education.

König, I. R., Fuchs, O., Hansen, G., von Mutius, E., and Kopp, M. V. (2017). What is precision medicine? European respiratory journal, 50(4).

Moriceau, G., Hugo, W., Hong, A., Shi, H., Kong, X., Clarissa, C. Y., Koya, R. C., Samatar, A. A., Khanlou, N., Braun, J., et al. (2015). Tunable-combinatorial mechanisms of acquired resistance limit the efficacy of braf/mek cotargeting but result in melanoma drug addiction. Cancer cell, 27(2):240–256.

Pribluda, A., de la Cruz, C. C., and Jackson, E. L. (2015). Intratumoral heterogeneity: From diversity comes resistanceheterogeneity and resistance. Clinical Cancer Research, 21(13):2916–2923.

Saunders, N. A., Simpson, F., Thompson, E. W., Hill, M. M., Endo-Munoz, L., Leggatt, G., Minchin, R. F., and Guminski, A. (2012). Role of intratumoural heterogeneity in cancer drug resistance: molecular and clinical perspectives. EMBO molecular medicine, 4(8):675–684.

Seifert, H., Fisher, R., Martin-Liberal, J., Edmonds, K., Hughes, P., Khabra, K., Gore, M., and Larkin, J. (2016). Prognostic markers and tumour growth kinetics in melanoma patients progressing on vemurafenib. Melanoma research, 26(2):138–144.

Silva, A. S., Kam, Y., Khin, Z. P., Minton, S. E., Gillies, R. J., and Gatenby, R. A. (2012). Evolutionary approaches to prolong progression-free survival in breast cancerprolonging progression-free survival in breast cancer. Cancer research, 72(24):6362–6370.

Smalley, I., Kim, E., Li, J., Spence, P., Wyatt, C. J., Eroglu, Z., Sondak, V. K., Messina, J. L., Babacan, N. A., Maria-Engler, S. S., et al. (2019). Leveraging transcriptional dynamics to improve braf inhibitor responses in melanoma. EBioMedicine, 48:178–190.

Strobl, M. A., West, J., Viossat, Y., Damaghi, M., Robertson-Tessi, M., Brown, J. S., Gatenby, R. A., Maini, P. K., and Anderson, A. R. (2021). Turnover modulates the need for a cost of resistance in adaptive therapyturnover modulates resistance costs in adaptive therapy. Cancer research, 81(4):1135–1147.

Suda, K., Tomizawa, K., Osada, H., Maehara, Y., Yatabe, Y., Sekido, Y., and Mitsudomi, T. (2012). Conversion from the “oncogene addiction” to “drug addiction” by intensive inhibition of the egfr and met in lung cancer with activating egfr mutation. Lung Cancer, 76(3):292–299.

Sun, C., Wang, L., Huang, S., Heynen, G. J., Prahallad, A., Robert, C., Haanen, J., Blank, C., Wesseling, J., Willems, S. M., et al. (2014). Reversible and adaptive resistance to braf (v600e) inhibition in melanoma. Nature, 508(7494):118–122.

Van Rossum, G. and Drake Jr, F. L. (1995). Python tutorial. Centrum voor Wiskunde en Informatica Amsterdam, The Netherlands.

Vasan, N., Baselga, J., and Hyman, D. M. (2019). A view on drug resistance in cancer. Nature, 575(7782):299–309.

Virtanen, P., Gommers, R., Oliphant, T. E., Haberland, M., Reddy, T., Cournapeau, D., Burovski, E., Peterson, P., Weckesser, W., Bright, J., van der Walt, S. J., Brett, M., Wilson, J., Jarrod Millman, K., Mayorov, N., Nelson, A. R. J., Jones, E., Kern, R., Larson, E., Carey, C., Polat, İ., Feng, Y., Moore, E. W., Vand erPlas, J., Laxalde, D., Perktold, J., Cimrman, R., Henriksen, I., Quintero, E. A., Harris, C. R., Archibald, A. M., Ribeiro, A. H., Pedregosa, F., van Mulbregt, P., and Contributors, S… (2020). SciPy 1.0: Fundamental Algorithms for Scientific Computing in Python. Nature Methods, 17:261–272.

Zhang, J., Cunningham, J. J., Brown, J. S., and Gatenby, R. A. (2017). Integrating evolutionary dynamics into treatment of metastatic castrate-resistant prostate cancer. Nature communications, 8(1):1816.

